# The phase of Theta oscillations modulates successful memory formation at encoding

**DOI:** 10.1101/2020.08.11.246421

**Authors:** Josephine Cruzat, Mireia Torralba, Manuela Ruzzoli, Alba Fernández, Gustavo Deco, Salvador Soto-Faraco

**Author notes:** Corresponding author: Josephine Cruzat, Universitat Pompeu Fabra, Mercè Rodorera Building, C/ Ramon Trias-Fargas, 25-27, 08005 Barcelona, Spain.

## Abstract

Several past studies have shown that attention and perception can depend upon the phase of ongoing neural oscillations at stimulus onset. Here, we extend this idea to the memory domain. We tested the hypothesis that ongoing fluctuations in neural activity have an impact on memory encoding using a picture paired-associates task to gauge episodic memory performance. Experiment 1 capitalized on the principle of phase reset. We tested if subsequent memory performance fluctuates rhythmically, time-locked to a reset cue presented before the to-be-remembered pairs. We found indication that behavioral performance was periodically and selectively modulated at theta frequency (∼4 Hz). In Experiment 2 we focused on prestimulus ongoing activity using scalp EEG recorded while participants performed the pair-associate task. We analyzed subsequent memory performance as a function of theta and alpha activity around the presentation of the to-be-remembered pairs. The results of the pre-registered analyses, using large electrode clusters and generic spectral ranges, returned null results of prestimulus phase-behavior correlation. However, we found that post-stimulus theta-power modulations in left frontal scalp predicted subsequent memory performance. This post-stimulus effect in theta power was used to guide a post-hoc prestimulus phase analysis, narrowed down to more precise scalp location and frequency. This analysis returned a correlation between prestimulus theta phase and subsequent memory. Altogether, these results suggest that the prestimulus theta activity at encoding has an impact on later memory performance.

## 1. Introduction

In our daily life we are often bombarded with myriad sensory stimuli and events of different kinds. Only a fraction of them are later remembered, whereas others are just forgotten. Although several processes influence our ability to remember, it is generally agreed that in all cases later recognition strongly relies on successful memory formation (Brassen et al. 2006; Wimber et al. 2010; Jutras et al. 2009). The subsequent memory paradigm has proven a successful procedure to study the neural underpinnings of memory formation in various types of memory systems (Paller et al. 1987; Hanslmayr and Staudigl 2014; for review see Paller and Wagner 2002). This paradigm usually compares neural activity that was registered following the onset of to-be-remembered stimuli (that is, at encoding), as a function of whether these stimuli are later remembered or not. Differences—known as subsequent memory effects (SMEs)—can help identify memory processes involved at encoding that facilitate later recall. Studies employing this approach have revealed that the brain activity during encoding is relevant for whether or not the stimulus will be subsequently remembered (Brewer et al. 1998; Wagner et al. 1998; Fernández et al. 1999; Kirchhoff et al. 2000; Strange et al. 2002; Sederberg et al. 2003; for review see Paller and Wagner 2002; Kim 2011). Many of these studies support the notion that oscillations in neural activity at the momnt of encoding, just after the stimulus is presented, are associated with effectiveness in later recall (for review see Klimesch 1999; Hanslmayr and Staudigl 2014). Evidence for the role of neural oscillations in episodic memory formation in humans comes from EEG/MEG studies using the SME approach. These studies show that increases in theta (4—8 Hz) and gamma (>30 Hz) power, and decreases in alpha (8—13 Hz) and beta (13—30 Hz) oscillatory power, are associated with more effective recall (Backus et al. 2016; Lega et al. 2012; Sederberg et al. 2003; Staudigl and Hanslmayr 2013; though see Sederberg et al. 2006, Greenberg et al. 2015 for counter examples of decreases in theta). While the association between oscillatory power and memory formation in humans has been investigated in numerous experiments, only a handful of studies have explored the potential role of the oscillatory phase. These studies provide evidence for a role of phase synchronization among different sensory cortices in the theta frequency during memory formation (Backus et al. 2016; Clouter et al. 2017).

The studies mentioned above have considered evoked or induced oscillatory activity after the stimulus to be encoded. In addition, previous results also suggest that ongoing fluctuations in neural activity even *before* stimulus presentation for encoding might also play a role in determining whether or not a stimulus will be later remembered (Park and Rugg 2010; Haque et al. 2015; Otten et al. 2006; Otten et al. 2010; Guderian et al. 2009; Fell et al. 2011; Rutishauser et al. 2010; Addante et al. 2015). Notably, the phase of the ongoing low-frequency oscillations appears to reset at the moment of the presentation of behaviorally relevant stimuli (Rizzuto et al. 2003; Haque et al. 2015). This shift in the dynamics of prestimulus ongoing oscillations might play a fundamental role in the synchronization of neuronal populations and the coordination of information transfer necessary for efficient encoding. In fact, some studies show that phase synchronization is more precise during encoding of information that is later remembered compared to later forgotten, most likely acting as the “gluing mechanism” for binding human memories (Buzsaki and Draguhn 2004; Backus et al. 2016; Hanslmayr et al. 2016; Clouter et al. 2017; for review see Fell and Axmacher 2011). Altogether, these findings suggest that memory formation not only depends on stimulus-driven processes but also on endogenous brain states at the time of stimulus presentation. Here, we set out to address this question: does the phase of prestimulus ongoing oscillations have an impact on memory formation? A similar question has been addressed by previous research in the domain of perception and attention in humans. In particular, some studies have related stimulus detection with the phase of the ongoing oscillations at or before stimulus onset. These studies have revealed cyclic alternations in behavioral performance mainly in the theta and alpha bands (∼4–8 and 8–12 Hz, respectively) (Busch et al. 2009; de Graaf et al. 2013; Klimesch et al. 2007; Mathewson et al. 2009; Palva and Palva 2007; VanRullen 2016; but see also Ruzzoli et al. 2019 for conflicting evidence). The main idea behind this phenomenon is that oscillatory neuronal activity reflects rhythmic fluctuations in the neuron’s membrane potential, which are associated with changes in neuronal excitability (Buzsaki and Draguhn 2004; Lakatos et al. 2005; Fries et al. 2007). These fluctuations can be reflected in the EEG signal when large neuronal populations are synchronized. In the case of perception, low-frequency oscillations may gate neural responses to incoming sensory information, producing peaks and troughs that correspond to favorable and unfavorable states for sensory processing. We hypothesize that if the phase of slow ongoing oscillations can modulate perceptual processing, then the phase of prestimulus fluctuations may also modulate encoding efficiency and hence, the success in later memory performance.

We tested this hypothesis using a paired-associates memory task in two experiments performed by separate groups of healthy human participants. Participants were instructed to memorize a short sequence of image pairs and then asked whether newly presented image pair probes had been previously associated or not.

In the first experiment, we used the logic of phase reset (Rizzuto et al. 2003) on behavioral performance. We tested if subsequent performance fluctuates rhythmically time-locked to a cue (phase-reset signal) presented before the to-be-remembered pairs of pictures. According to our hypotheses, we focused the analysis on low frequencies, ranging from 2 to 20 Hz. In the second experiment, we tested the role of ongoing low-frequency neural oscillations—before the presentation of the picture pairs—in subsequent recognition performance using EEG. Specifically, we focused on the phase and amplitude of alpha and theta oscillations. To do so, we examined prestimulus changes in the EEG as participants performed the task adapted from Experiment 1 but capitalizing on spontaneous neural oscillations, instead of those arising from phase reset. The hypotheses and analysis pipeline of Experiment 2 were pre-registered based on the results of Experiment 1 (see below; pre-registration available at https://osf.io/4f5qc/). To anticipate the results, although the pre-registered analysis pipeline did not yield significant results, we found and report phase-behavior correlations upon further exploration of the data following a data-driven approach.

## 2. Methods Experiment 1

### 2.1 Ethics Statement and Participants

The Clinical Research Ethical Committee of the Municipal Institute of Health Care (CIEC-IMAS) Barcelona, Spain, approved the study. Following the Declaration of Helsinki, all subjects gave written consent before their participation.

Data from 30 healthy subjects (21 females, mean age 23.2 ± 4.5 years, 5 left-handed) were used in the behavioral study. Data from 8 additional subjects were excluded based on the predefined inclusion criteria (demographic characteristics of each subject are detailed in supplement Table S1). The inclusion criteria were set up to ensure both, a sufficient number of trials in each response category (hit/miss), and that participants responded above chance level (for more details about inclusion criteria see Experimental Design and Procedure). Participants were compensated with 10€/hour.

### 2.2 Experimental Design and Procedure

Participants performed a visual paired-associates memory task (Figure 1A) adapted from Haque and colleagues (Haque et al. 2015). In our version of the paradigm, participants were asked to memorize pairs of pictures instead of pairs of words. Participants sat ∼60 cm away from a 21-inch CRT computer monitor (60 Hz refresh rate). Stimulus presentation was controlled using Matlab (Version R2016a, The MathWorks, Inc., MA, USA) and the Psychophysics Toolbox (Brainard 1997; Kleiner et al. 2007).

**Figure 1.**
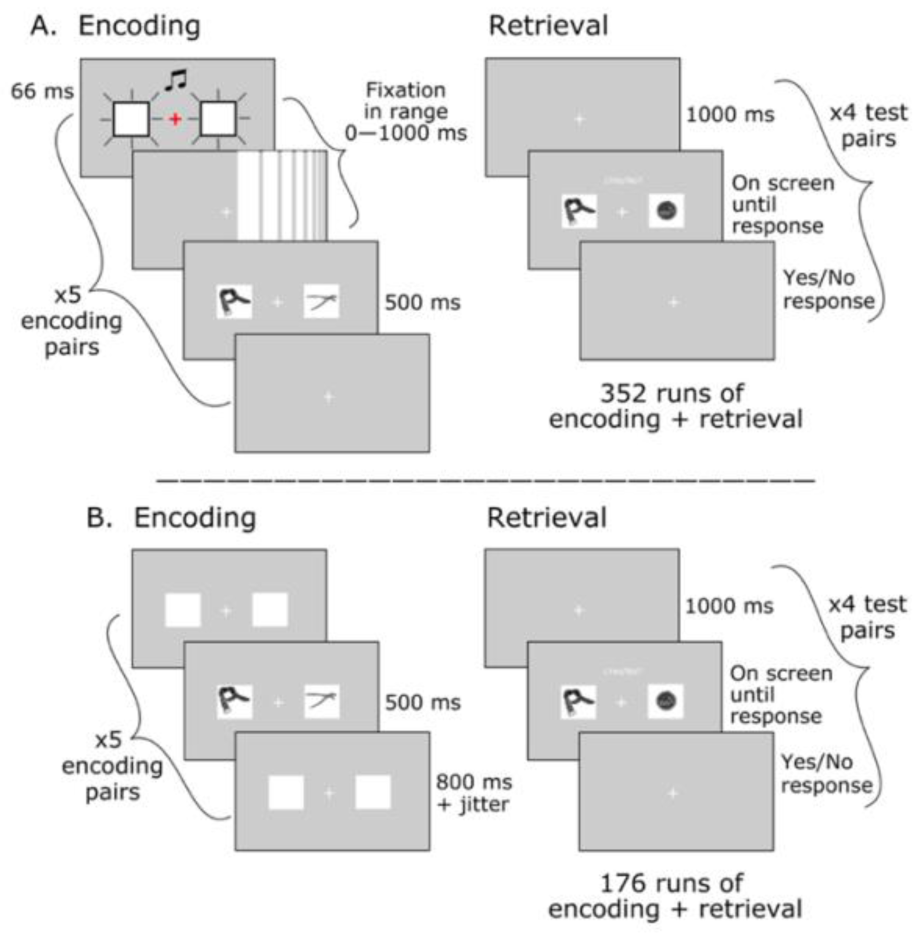
Experimental design. A, Experiment 1. Participants performed a visual pair-associates memory task. During the encoding block, participants were asked to learn five unrelated image pairs presented side-by-side on placeholders for 500 ms. A cue—composed by a central fixation cross and placeholders—flashed once synchronously together with a sound-beep before each image pair presentation. Critically, the time-lag between the cue and the image pair to be encoded (the cue-to-target interval) was varied randomly between 0 and 1000 ms. Each encoding block was followed by a four-trials recognition block where participants judged whether a given image pair had been presented together in the previous encoding block. Each participant provided a total of 1.408 responses. B, Experiment 2. The experimental design was identical to Experiment 1 with the following exceptions: During the encoding block, image pairs were sequentially presented at a variable stimulus-onset asynchrony (SOA), with a jitter according to an exponential distribution (mean 1s) to compensate for hazard rate and prevent temporal expectation. Each image pair was followed by a blank inter stimulus interval (ISI) of 800 ms. Each participant performed half of the task, thus providing a total of 704 responses.

Due to the large number of trials, the experiment was completed over two sessions on different days within the same week (on average, three days apart). In total, it consisted of 352 runs, each composed of an encoding block of five pairs followed by a recognition block with four memory probes. Each participant provided a total of 1.408 (352×4) responses throughout the experiment. During the encoding blocks, five image pairs in placeholders were presented in sequence with a central fixation cross in between them (placeholders and fixation were present throughout the block). Before each image pair, an audio-visual reset cue was presented. The reset cue consisted of a flash of fixation and placeholders synchronized with a sound and was intended to modulate cortical excitability by phase resetting the ongoing oscillatory activity (Lakatos et al. 2009; Fiebelkorn et al. 2011; Daitch et al. 2013). Critically, the interval between the reset cue and the target image pair to be encoded (cue-to-target interval) varied for each trial within each run between 0 and 1000 ms in steps of 16.66 ms (leading to 61 unique times with ∼44 trials per interval in the (0 to 500 ms) window of interest, and ∼16 trials from 500 ms onwards). Then, each image pair appeared for 500 ms, followed by 1500 ms interval that led to the next reset cue. We adopted this strategy to appreciate potential oscillations in behavior without using EEG to measure ongoing brain signal fluctuations. Since, based on the hypothesis, we focus on fluctuations >2 Hz, our window of interest was 0 to 500 ms. Hence, the probability of occurrence of targets within the 0-500 ms interval was about three times that of long-time intervals (500 to 1000 ms). Long-time intervals were only included to make the stimulus onset less predictable within the times of interest, but they were not used for the analysis. We expected the audio-visual cue to reset the phase of the ongoing oscillations, so then behavioral performance at the recognition blocks should present an oscillatory pattern time-locked to the cue presentation.

Each encoding block was followed by a recognition block that consisted of a sequence of four test image pairs. On each test pair, participants had to judge whether the two images had been presented together (paired) in the preceding encoding block. We tested four pairs as a test of all five pairs would make the fifth pair response predictable. The test pairs always contained items that had previously appeared in the encoding block (whether paired or not), so that the task could not be solved by just recognizing the appearance of new items. Each image pair was preceded by a prompt—a row of question marks—that appeared on the screen for 500 ms followed by a blank interval of 1000 ms. The image pair, placeholders, and fixation cross remained on the screen until the response (yes/no) was collected through the keyboard. To minimize the number of false alarms, participants were encouraged to respond yes only when they were confident about their response and to say no if doubting. Image pairs in the recognition block could be a match (if they had appeared together in the encoding block) or a mismatch (if they appeared in different pairs during encoding). Matched pairs were slightly more likely (60% on average), compared to mismatch pairs, to ensure enough useful trials for analysis. From a total of 1.408, on average, 847 were match trials (min 831, max 865). Each recognition block could have a different number of yes/no correct responses to prevent guessing based on response history. After every 10 runs (encoding plus recognition blocks), participants took a break and were updated on their overall performance to control for false alarms and keep them engaged in the task.

Prior to analysis, we established a set of individual inclusion criteria regarding performance: (1) false alarm rate below 15%, (2) hit rate above 35% to ensure that with the maximum false alarms rate accepted the participants responded above chance level, and (3) hit rate below 80% to avoid ceiling effects. These criteria were defined to ensure enough number of trials in each response category (hit, miss), allowing the possibility to capture fluctuations in behavior—in the case they existed.

### 2.3 Stimuli

The pictures used in the pairs were images of familiar objects in a usual perspective, from a wide range of semantic categories selected from two databases: The Bank of Standardized Stimuli (BOSS) (Brodeur et al. 2010; Brodeur et al. 2014) and the set of 2400 Unique Objects used in Brady et al. (2008) (Brady et al. 2008). Stimulus characteristics are described in detail in previous studies (Brodeur et al. 2010; Brodeur et al. 2014; Brady et al. 2008). The two sessions of the experiment used different image sets. Within a pair, images belonged to different semantic categories. Living objects were not mixed with non-living objects either on the same trial or block (80% of the pairs corresponded to non-living objects). All images were converted to black and white to avoid memory facilitation based on color associations (Lewis et al. 2013). Image size was subtending a square of approximately 5 degrees of visual angle. The auditory stimulus used as resetting signal was a beep sound (4000 Hz, ∼60 dB, 66 ms) presented through headphones.

### 2.4 Behavioral Analyses

The goal in this experiment was to address if behavioral performance at recognition fluctuated rhythmically as a function of the reset cue (cue-to-target interval) at encoding. We followed the analytical method described in Fiebelkorn et al. (2013) to extract rhythmic variations of behavioral performance. For each participant, we only used responses to match trials to obtain hit and miss rates. To get a temporally smoothed series relating memory behavior to encoding time, we first calculated the hit rate in a sliding 50 ms window (16.66 ms steps) for the whole window of interest (from 0 to 500 ms), and we averaged time-dependent hit rate across participants. Then, we used a Fast Fourier Transform (FFT) to extract the spectral pattern of the detrended behavioral time series and focused our analyses in the 2–20 Hz frequency range. The minimum and maximum frequencies of interest were determined by the window of interest (500 ms) and the width of the sliding window used to calculate time resolved hit rate (50 ms). We sought for oscillatory patterns above and beyond chance within the low-frequency spectral window.

To examine the reliability of the results, we assessed statistical significance using a non-parametric procedure with 10.000 randomly generated surrogates. For each participant, we permuted hit rates across the cue-to-target interval windows, averaged across participants, and extracted the surrogate power spectrum. The *p-*value corresponds to the proportion of permutations in which the surrogate value matched or exceeded the empirical value. We obtained the *p-*values for each frequency (9 frequencies) and then applied multiple comparisons correction with False Discovery Rate (FDR) using the Benjamini-Hochberg procedure (Benjamini and Hochberg, 1995) with an alpha level of 0.05.

## 3. Results Experiment 1

We expected the time-resolved recognition performance to show a periodic component indicative of fluctuations time-locked to the cue at the encoding. Overall, participants performed above chance level with a mean hit rate of 62.97 ± 10.74%, and a false alarm rate of 6.09 ± 3.82% (mean hits 534, min 323, max 672; mean misses 314, min 169, max 516; individual hit and miss rates plus demographic data are detailed in the supplemental Table S1). In line with the question of this first experiment, behavioral performance in recognition fluctuated as a function of cue-to-target interval (Figure 2A). We used FFT on the behavioral time series and found that memory performance was periodically and selectively modulated with a peak frequency of ∼4 Hz (Figure 2B). Although this result was statistically significant (*p*=0.0398), it did not survive correction for multiple comparisons across all frequencies included in the test (2–20 Hz).

**Figure 2.**
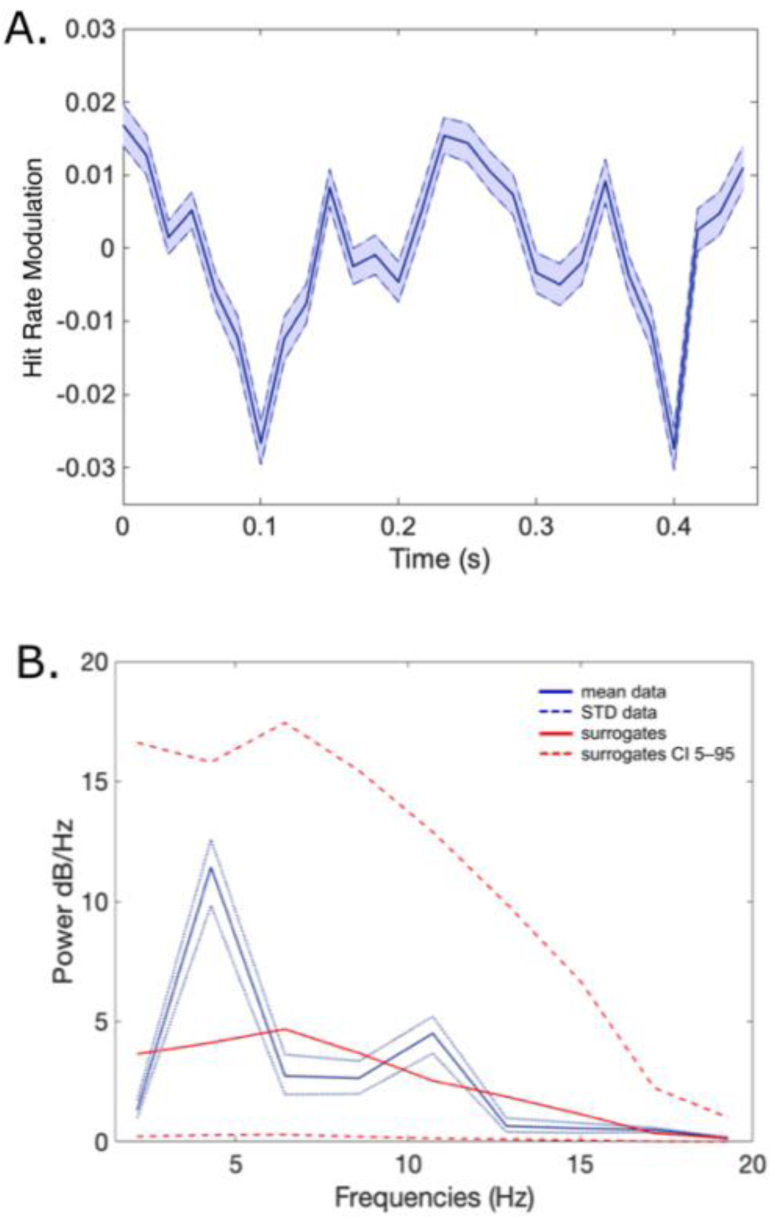
A. Behavioral Performance. Demeaned mean hit rate and standard deviation across participants as a function of time (N=30). **B. Power Spectrum**. Amplitude measurement obtained using the FFT for the mean hit rate. As can be seen (blue line), there is a peak around ∼4 Hz (theta band).

Because of the large spectral window analyzed—and the ensuing strong correction for multiple comparisons—our data provide only weak support for the hypothesis of a rhythmic modulation of behavioral performance within the theta range, after phase resetting by the prestimulus cue. This putative modulation, if only tentatively, suggests that the phase in the theta band at which the stimulus arrives has an impact on later successful recognition. Despite the relatively large spectral window analyzed and the unknown statistical power of the effect, we considered this result potentially indicative of the relevant frequency, and hence useful to narrow down on a more specific spectral window of interest for an EEG study, presented in Experiment 2.

## 4. Methods Experiment 2

The aim in Experiment 2 was to investigate the role of ongoing low-frequency oscillations—before the to-be-encoded stimulus presentation—in subsequent recognition performance using EEG. Several neuroimaging studies have shown that frontotemporal regions are actively engaged during episodic encoding (Wagner 1999; Kirchhoff et al. 2000; Hanslmayr et al. 2011; Park and Rugg 2011; Griffiths et al. 2016). However, the results show contradictory evidence for theta frequency fluctuations of the local field potential: some provide evidence in favor of encoding-related increases (Herweg et al. 2016; Addante et al. 2011; Summerfield and Mangels 2005; Osipova et al. 2006; Hanslmayr et al. 2011), other results report decreases (Fellner et al. 2016; 2019; Michelmann et al. 2018). A well-accepted interpretation of these activations is that they reflect hippocampal theta, induced in cortical areas via hippocampal-cortical feedback connections (for a review see Herweg et al. 2020). Moreover, episodic-encoding is also associated with decreases in the alpha band in occipitoparietal regions (Klimesch and Doppelmayr 1996; Klimesch et al. 1997; 2000), likely contributing to the encoding of visuospatial stimulus attributes and signaling directed attention (Klimesch 1999; Klimesch et al. 2000). On these grounds, we focused our analysis on the phase and amplitude of the theta and alpha oscillations in broad frontotemporal and occipitoparietal regions, respectively. Here, we used the 4–7 Hz range to clearly distinguish between theta and alpha activity (which is commonly defined as 8–12 Hz). Please note that although theta is usually defined as the 4–8 Hz range, the 4–7 Hz range has also been used (Crivelli-Decker et al. 2018; Hanslmayr et al. 2010; Li et al. 2017; Mizrak et al. 2018).

We hypothesized that the phase of the ongoing oscillations at which the to-be-encoded stimulus arrives modulates subsequent visual memory performance. If so, we expected to find fluctuations in behavioral performance as a function of the phase angle of the ongoing brain oscillation. The hypotheses and the initial analysis pipeline of this experiment were pre-registered (https://osf.io/4f5qc/). In the results section below we indicate which part of the analyses belong to the pre-registered pipeline, and which ones are exploratory.

### 4.1 Participants and Ethics Statement

As in Experiment 1, the local ethics committee approved the study, and all participants gave written consent before their participation.

Data from 30 subjects (15 females, mean age 22.9 ± 2.9 years, 5 left-handed) were used in the EEG study. Data from six additional subjects were excluded: 2 did not meet the behavioral inclusion criteria specified above, and 5 because of artifact contamination of the EEG (demographic characteristics of each subject are detailed in supplement Table S2). Participants were compensated with 10€/hour.

### 4.2 Experimental Design and Procedure

The experimental design and procedure were identical to Experiment 1 with the following three exceptions: First and foremost, since ongoing oscillations were registered directly with EEG, we did not use a reset cue. During the encoding blocks, image pairs were sequentially presented at a variable inter-stimulus interval (ISI), without a prestimulus cue. Indeed, we expected the image pair presentation to work as a reset cue and link the phase of the ongoing oscillation at the image pair onset with subsequent behavioral recognition. The second difference was that the ISI was composed of a fixed 500 ms period plus a jitter according to an exponential distribution—with mean 1000 ms—to further increase temporal uncertainty (Figure 1B). And third, participants performed only one experimental session containing a total of 704 responses from which, on average, 421 were match trials (min 412, max 431). The reduction in observations was justified by the advantage of measuring multiple frequency bands within one trial through EEG, and by the possibility to narrow down on the frequency band of interest for the analysis according to the result in Experiment 1 and hypothesis.

### 4.3 Stimuli

The stimuli were identical to the ones used in the first session of Experiment 1, namely, images taken from the Bank of Standardized Stimuli (BOSS) (Brodeur et al. 2010; Brodeur et al. 2014). All the stimuli were processed as described for Experiment 1.

### 4.4 EEG Recording and Data Analyses

Electrical brain signals were recorded using an EEG system with 60 active electrodes (actiCAP, Brain Products GmbH) located according to the standard international 10–10 system and sampled at a rate of 500 Hz. Two electrodes placed at the right and left mastoids served for offline re-reference. An electrode placed at the tip of the nose served for online reference, and the ground electrode was placed at AFz. Electrooculogram (EOG) was monitored on the horizontal and vertical directions to control for blinks and eye movements.

All data and statistical analyses were performed using custom code in Matlab (Version R2016b, The MathWorks, Inc., MA, USA), and the FieldTrip toolbox (Oostenveld et al. 2011). Raw EEG data were re-referenced to averaged mastoids, and a notch-filter at 50 Hz was applied to remove line contamination. To extract specific spectral information, we bandpass filtered the EEG data in two preselected frequency bands: theta (4–7 Hz) and alpha (8–14 Hz), using a second-order zero-phase Butterworth filter. The filtered data were then segmented from –500 ms to 100 ms with respect to the onset of each pair in the encoding blocks. Malfunctioning electrodes were removed, and their data were estimated based on neighboring electrodes using spline interpolation. Trials contaminated by blinks, muscle movements, or brief amplifier saturation were discarded from further analyses by visual inspection. As in Experiment 1, we were interested in the encoding phase of trials labeled as hit or miss according to participant’s response in the subsequent recognition block. Thus, only matching trials could be used for the analysis. To ensure a reliable estimation of oscillatory patterns, for a given participant, if the number of artifact-free trials in the less populated condition (hit or miss) was below 80, the entire participant’s dataset was excluded.

Since we were interested in phase-dependent memory effects, our hypothesis capitalized on the endogenous oscillations in the EEG before the onset of image pairs at the encoding blocks. We compared the spectral power and the instantaneous EEG phases between hits and misses. Based on the results of Experiment 1 and previous literature (Addante et al. 2011; Sederberg et al. 2003; Nenert et al. 2012; Jensen et al. 2002; Schack et al. 2002; Summerfield and Mangels 2005), we set the spectral-spatial regions of interest (ssROIs). The frequencies and regions of interest were theta (4–7 Hz) for frontotemporal electrodes, and alpha (8–14 Hz) for occipitoparietal electrodes (see details in Figure 3). Phase opposition measurement values at electrodes belonging to the ssROIs at the specific frequency of interest were averaged at the individual participant level and then averaged over participants.

**Figure 3.**
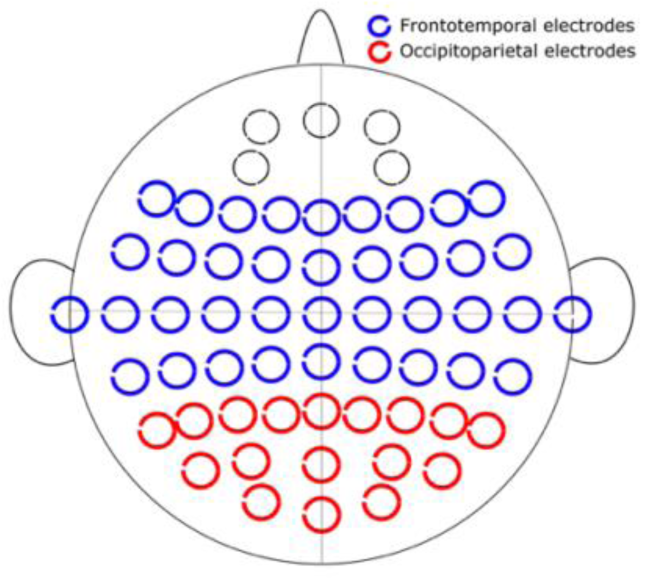
Map of scalp EEG electrode locations and their corresponding regions of interest (ROIs). Electrodes were divided into 2 ROIs: frontotemporal—outlined in blue, and occipitoparietal—outlined in red.

The Hilbert transform was used on the narrow-band filtered data to obtain, for each frequency at time, the associated complex analytical signal (*t*), defined by the instantaneous phase *φ*(*t*)We then calculated the intertrial coherence (ITC) separately for hits and misses for the time interval -500 to 100 ms relative to the image-pairs onset. The ITC is a direct measure of frequency-specific synchronization (Lachaux et al. 1999), and is given by,

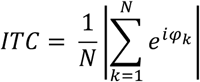

Where *N* is the total number of trials and k the trial ID. Under complete independence across trials between the EEG data and the time-locking events, ITC is nearly 0, representing absence of synchronization; whereas for ITC equals 1, all phases are concentrated, representing full synchronization.

#### Phase Opposition Analyses

To determine subsequent memory effects as a function of the phase at stimulus onset, we evaluated phase differences between hits and misses using the Phase Opposition Sum (POS) (VanRullen 2016), see pre-registration (https://osf.io/4f5qc/). The POS measurement estimates the correlation between an oscillation phase at a particular frequency, and an observed behavior. It is computed as the sum of the ITC calculated separately for two or more conditions according to the observed behavior. Moreover, it assumes that the phase in the prestimulus period is distributed randomly (i.e., follows a uniform distribution) across trials and, consequently, if the EEG phase at one point in time and frequency influences subsequent memory recognition, we expect a phase concentration opposition between hits and misses. Analytically, the POS measure is given by:

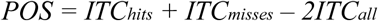

where we computed the ITC individually for hits and misses.

According to the simulation results presented in VanRullen (2016), an imbalanced number of trials between conditions decreases the sensitivity of the index. Based on the performances obtained in Experiment 1, we expected an imbalance in the number of trials in each category (number of hits larger than misses). In consequence, we decided to equate the number of trials between conditions for analysis by subsampling the more populated condition. By equating the number of trials using subsampling, we estimated that the POS per subject would be based on approximately 300 trials, which according to VanRullen (2016), provides around 80% statistical power for a balanced number of trials. For each subject, we calculated the ITC of the less populated condition using all the trials available for this condition (N). Subsequently, N trials were randomly sampled from the more populated condition and the ITC for this condition and for both conditions (ITC_all_) were calculated. We repeated the calculation of the ITC for both the more populated and the collapsed conditions together 250 times and we averaged across repetitions to obtain an estimation of the ITC of the more populated condition and both conditions together and we used these estimations to calculate POS. We averaged the POS values across electrodes within each region of interest (ROI).

The POS is bounded between -1 and 1. If the phase is related to the trial outcomes, then the POS comes out positive (ITC of each trial group should exceed the overall ITC), then the *p*-value corresponds to the proportion of group average pseudo-POS larger than the actual POS. To test for significance, we used the Monte Carlo test procedure i.e. for each subject, we created a distribution of 500 surrogates by randomly assigning the hit/miss labels to existing trials. These surrogate individual distributions were subsequently used to calculate 10,000 group average pseudo-POS by randomly picking surrogates of the distribution of each participant. *p*-values were further corrected for multiple comparisons using the False Discovery Rate (FDR) method (p<0.05).

In the post-hoc analyses that we present after the pre-registered ones, we used an additional measure for phase opposition called the Phase Consistency Metric (PCM) (Landau et al. 2015). Similar to the POS method, the PCM quantifies the consistency of phase differences in a particular frequency band but with a bias-free sample estimator that controls for the imbalance in the number of trials between conditions by looking at pairs of observations instead of all observations together. The PCM evaluates the average of the cosine of the differences between all possible pairs of hit-miss trials. In case of perfect opposition, the phases would be point to 180° apart and, therefore the observed value would be -1. Therefore, negative values in PCM would indicate phase opposition.

## 5. Results Experiment 2

Participants performed the task with an overall hit rate of 64.15 ± 6.89% and a false alarm rate of 7.46 ± 3.52% (mean hits 270, min 190, max 321; mean misses 151, min 98, max 222; see individual results in supplemental Table S2). After artefact rejection, the number of trials per participant was 343 ± 32 trials (minimum 280, maximum 401), so 79 ± 32 trials per subject were discarded (minimum 21, maximum 142). The focus of this experiment was to address whether the phase of the ongoing EEG activity before the associate-pair onset at encoding (−500 to 0 ms time-window) could predict subsequent memory performance.

To answer this question, according to the pre-registered analysis pipeline, we used the POS analysis (VanRullen, 2016) described above, on the preselected frequencies and regions of interest. The POS results showed no significant phase opposition in either Theta (max 0.044 ± 0.017, t=-500 ms) or Alpha frequency bands (max 0.0509 ± 0.026, t=-174 ms) in either ROI (Supplemental Figure S1, A-B). In principle, this result would mean that the phase of ongoing oscillations within the specified frequency ranges and ROIs was not predictive of subsequent memory performance (or that if present, the effect was undetectable with the present statistical power). However, before committing to this conclusion, we explored the data further to rule out potential alternative explanations for the null result from the pre-registered analysis and to try and increase the sensitivity of the measurements. The results of the exploratory analyses are reported below.

### 5.1 Reality checks and alternative phase-opposition analysis

First, an important reality check is to ensure the assumption that the phase of ongoing brain activity before the stimulus in the time period of interest is distributed randomly. This ensures that ongoing phase is not influenced by any stimulus-related factor or anticipation based on the protocol itself. As expected, through the calculation of phase concentration with the Rayleigh statistic, we observed that prestimulus phases were randomly distributed when collapsing hits and misses in the prestimulus time-window that corresponded to pure ongoing activity. We observed phase concentration around stimulus onset time (∼1 cycle of each frequency of interest), which is due to the use of non-causal filters, and the stimulus-related phase resetting.

Second, we questioned whether the Phase Opposition Sum (POS) index used in the main analyses was reliable within the range of parameters we used. As it was explained above, the POS metric has a positive bias (inherited from the ITC used in the calculation) that correlates with the total number of trials and with the relative number of trials per condition (VanRullen 2016; Moratti et al. 2007; Vinck et al. 2010). It has been shown that the statistical power of POS and other phase opposition measures decreases as the imbalance in trials between conditions increases (VanRullen 2016). Our EEG experiment had two sources of imbalance in the number of trials for the hit vs. miss comparison: (1) because conditions were determined as a function of performance, we could not anticipate exactly how many trials per condition there would be, and (2) due to the unequal artifact rejection. For this reason, in the planned analyses above we equated the number of trials in each condition by random picking from the more populated condition the same number of trials as in the less populated condition. This procedure, however, may have been suboptimal, as it resulted in a decrease in the total number of trials available to estimate each of the POS in each subject.

Additionally, the number of repetitions used to calculate POS was based on computational time. In order to evaluate if the number of repetitions used to calculate POS was enough to produce a consistent POS, we calculated the empirical POS and generated null POS measures for one electrode and time point in the theta frequency band, for different numbers of random samples (from 25 to 1.000 samples). We observed that even when using 1.000 random samples per participant, the *p-*values obtained were not stable: at the maximum number of iterations, the p-values varied in the interval 0.06 to 0.12 (Supplemental Figure S2). This result casts doubts on the stability of the POS obtained by the subsampling of the data and hence, on the null result observed.

Third, because of the above, we decided to ascertain whether phase effects could be found in our data when using an alternative measure, less affected by trial imbalance. We assessed the impact of the phase of ongoing EEG at encoding onset on subsequent performance using the Phase Consistency Metric (PCM) (Landau et al. 2015) method. Similar to the POS, the PCM quantifies the consistency of phase differences in a particular frequency band but with a bias-free sample estimator that controls for the imbalance in the number of trials between conditions by pairing observations. Like the POS, we did not observe any significant phase opposition in any frequency or region of interest when comparing the PCM between hit and miss trials (theta in frontotemporal electrodes: min 0.00013 ± 0.0022, t=-500 ms; alpha in occipitoparietal electrodes: min: 0.000087 ± 0.0027, t=-350 ms) (Supplemental Figure S1, C-D).

Fourth, we investigated whether our data could replicate previous results showing that differences in oscillatory amplitude in the time window around stimulus onset at encoding are predictive of subsequent memory performance (Strunk and Duarte 2019). In order to obtain time-resolved data, oscillatory amplitude values were computed using the Hilbert transform (as specified in the methods section) on a trial by trial basis for theta and alpha frequencies. Power was averaged across electrodes within each ROI. The difference between conditions in the period – 500 to 100 ms relative to stimulus onset, was estimated as follows (power expressed in dB):

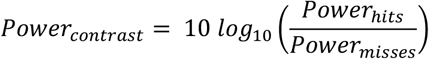

We assessed the significance of the power contrast between hits and misses within ROIs using a two-tail *t-test* corrected for multiple comparisons using the FDR method (p<0.05). In the alpha band, no significant differences in power between hits and misses were found (*t-test*, p<0.05, FDR corrected). However, we found a significantly higher theta power for later remembered stimuli compared to latter forgotten in the peri-stimulus time-window -50 to +100 ms (*t-test*, p<0.05, FDR corrected) (see, Figure 4). This subsequent memory effect (SME), even if including a brief prestimulus period, is most likely fully explained by evoked activity, given the temporal smoothing involved in the analysis used. SME in theta evoked activity is well in line with previous findings showing that increases in theta oscillatory power during the encoding period predict subsequent recall (Guderian et al. 2009; Sederberg et al. 2003; Osipova et al. 2006; White et al. 2013; Long et al. 2014; Solomon et al. 2019).

**Figure 4.**
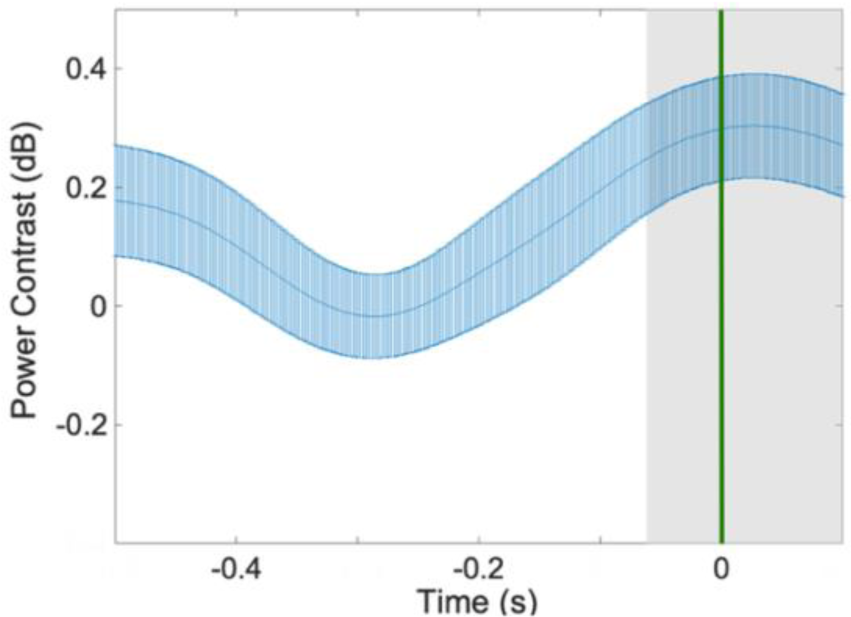
Theta power contrast between hits and misses (thick blue line). The Shaded grey area shows the period of significant differences (p<0.05) after FDR correction. The shaded blue area represents the standard error of the mean (SEM), and the green line shows stimulus onset.

### 5.2 Post-hoc analyses of phase effects using fine-tuned frequency and region of interest

Based on prior evidence for SMEs in evoked theta power (Backus et al. 2016; Lega, Jacobs, and Kahana 2012; Sederberg et al. 2003; Staudigl and Hanslmayr 2013), and on the significant theta power contrast for hits vs. misses in Experiment 2, we decided to adopt a data-driven approach to further seek for prestimulus phase effects using fine-tuned parameters.

Theta power modulations following stimulus presentation have been widely related to subsequent memory performance. Whereas several studies support that increases in theta amplitude during the encoding period relate to successful memory performance (Osipova et al. 2006; Sederberg et al. 2003; White et al. 2013; Clouter et al. 2017; Khader et al. 2010; Guderian et al. 2009; Long et al. 2014; Lega et al. Kahana 2012), others have reported the effect in the opposite direction (Long et al. 2014; Lega et al. 2012; Greenberg et al. 2015; Sederberg et al. 2006). Regardless of the directionality of the theta power effect, which could respond to two different mechanisms that arise from cortical and subcortical processes in support of memory encoding (Herweg et al. 2020), we decided to tailor the new pre-stimulus phase analysis to the frequency peak and scalp region based on this theta power SME. This approach is justified given that the a priori ROIs we used were very broad and probably suboptimal. Moreover, individual theta frequency shows large inter-individual differences (Haegens et al. 2014; Klimesch 1999), as it covaries with the individual alpha frequency (Doppelmayr et al. 1998; Klimesch and Doppelmayr 1996). Below, we describe the fine-tuning of the frequency of interest and scalp location adopting a data-driven approach. Later, we report the results of the ensuing prestimulus phase SME

#### Adjustment of Individual Frequency of Interest (IFOI)

To select the individual frequency of interest (IFOI) we calculated the scalp average power spectrum during the encoding period (0 to 1000 ms from stimulus presentation) using a Fourier Transform (FT) (padded to 12 s to increase frequency resolution, 4 slepian tapers) in the range from 1 to 40 Hz. Then, we calculated the power spectrum for all trials (hits and misses), and we normalized with respect to the mean power across frequencies (*P*_*m*_ = ⟨*P*(*f*)⟩_*f*_).:

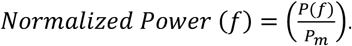

The IFOI was defined as the largest local maxima in the 3 to 7 Hz interval. We shifted the lower bound of the spectral range of the IFOI to account for participant variability. To illustrate the results, Figure 5 shows the scalp average power spectrum during the encoding interval for a representative participant. The mean IFOI measured was 4.05 ± 0.27 Hz (minimum 4 Hz, maximum 5.5 Hz).

**Figure 5.**
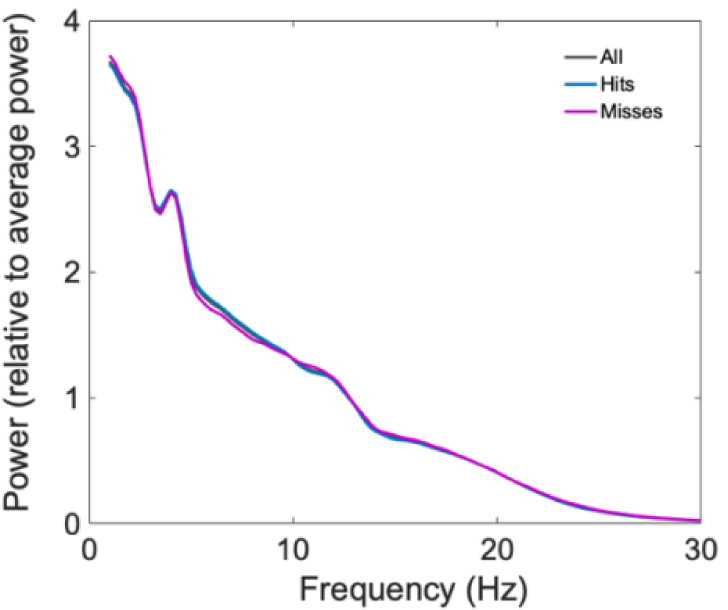
Individual frequency of interest (IFOI) for one representative participant. The figure represents the scalp average power spectrum, relative to average power, during the encoding interval for the collapsed distribution of hits and misses (black), as well as separately for hits (blue) and misses (red).

#### Adjustment of the Region of Interest (ROI)

We band-passed the signal around the individual frequency of each participant in the narrow band IFOI ± 2 Hz using a second-order Butterworth filter, and computed the corresponding instantaneous power spectrum using the Hilbert Transform. Then, the power contrast for hit vs. miss trials was obtained for each participant, electrode, and latency (0 to 500 ms in steps of 2 ms) and transformed to dB. As shown in Figure 6, across subjects, we identified a significant positive time-frequency cluster (*p-*value*<0*.*0002*) that initiated in the left frontal area—shortly after stimulus presentation—and extended towards central and posterior regions. Consequently, the new ROI was defined as the set of electrodes in the left frontal area which first displayed a significant power effect in the post-stimulus period (0-50 ms) and remained significant for most of the time within the window of analysis (0-500 ms): F3, F5, F7, FT7 and FC5. Significance in power across conditions was obtained by means of a *t-test* (right-tail, alpha level p=0.05) and corrected for multiple comparisons using a cluster approach (Maris and Oostenveld 2007), (alpha level p=0.05, 10.000 randomizations).

**Figure 6.**
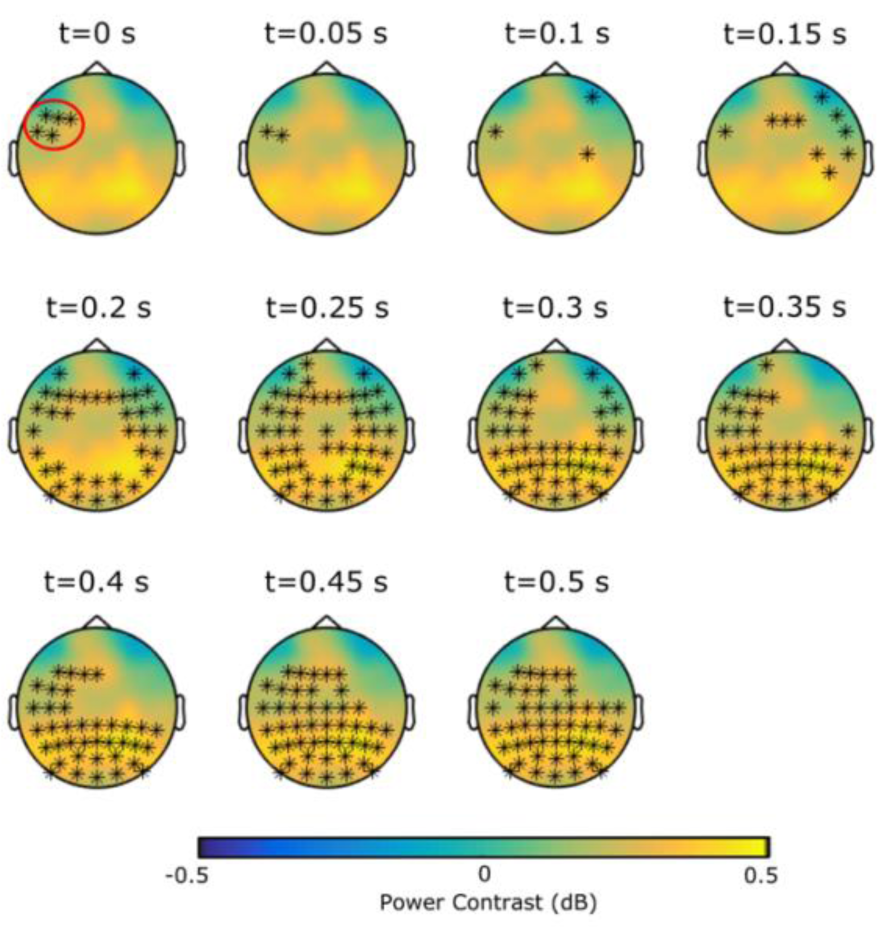
Adjusting the ROI based on the IFOI. Snapshots of the positive cluster for theta power contrast between hit and miss trials, using the individual frequency of interest (IFOI). Black stars indicate electrodes with significant activity at displayed time points. Red circle indicates the new ROI that was defined as the set of electrodes in the left frontal area which first displayed a significant power effect in the post-stimulus period.

#### Fine-tuned analysis of prestimulus phase SME

We ran the Phase Consistency Metric (PCM) analysis in the prestimulus time-window for hits vs. misses with the new, fine-tuned IFOI and ROI estimated as described above. The PCM allows to include all available trials without inducing a bias. For each participant, phases were extracted from the IFOI-filtered signal using the Hilbert Transform in the time-window immediately before stimulus onset (−500 to 0 ms), and the PCM was computed for each time point of the window. Because the surrogate distributions used for assessing the POS and PCM seemed to be biased by the data (as can be seen in Supplemental Figure S1), statistical significance for PCM values below the null hypothesis was assessed by means of surrogate distribution in the time domain.

The surrogate distributions were calculated for each subject and electrode as follows: For each trial, we randomly assigned the labels hit and miss, and then picked the phase of the EEG signal at a time point selected randomly from the -800 to -500 ms pre-stimulus window. We repeated this procedure 500 times per subject and electrode, and then averaged across electrodes. The individual null distributions were used to generate 10.000 group average null PCMs. We observed that PCM was not significantly below the null distribution (all p-values>0.5). Therefore, our results do not support phase opposition. But it could still be possible that either hit or miss trials were concentrated around a preferent phase, even if not opposite. We performed this final check, evaluating phase concentration of hit (later remembered) and miss (later forgotten) trials separately, by means of pairwise phase consistency (PPC) (Vinck et al. 2010), according to the following formula:

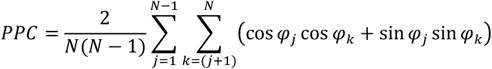

Where *N* is the total number of trials, and *φ*_*j*_ corresponds to the phase of trial j. The PPC is a measure of phase concentration that can take values between -1 and 1, that has been shown to be less biased by the number of trials than ITC (Vinck et al. 2010). When phases are perfectly aligned, the PPC equals 1.

PPC was calculated for the time window of interest, and all the electrodes of the ROI, for the theta filtered signal. The PPC for hit trials (PPC_Hits_) and for miss trials (PPC_Misses_) were calculated separately and statistical significance was assessed by means of a Montecarlo permutation test. Null distributions for hit and miss trials were built same as described above (label and time shuffled), with the number of hits and misses equivalent to the empirical values of hits and misses for each subject. After averaging across electrodes, we obtained for each subject a distribution of 500 null PPCs for hits and 500 null PPCs for misses. These individual PPCs were used to generate 10.000 group averaged null PCCs for hits and misses. The p-value corresponded to the proportion of times that the null distribution was above the empirical PPCs. We used Guthrie and Buchwald correction for correcting for multiple comparison (estimated autocorrelation 0.999, N=30, 251 time points, required minimum number of consecutive significant samples=26).

Around stimulus presentation onset, both hit and miss trials were significantly concentrated (Figure 7). This can be attributed to the phase resetting caused by the stimulus presentation. Miss trials were concentrated from -356 ms to 0 ms, but not in the early time window. On the other hand, hit trials were significantly concentrated for all the time window of interest -500 to 0 ms (Figure 8). In other words, the results show that there is an increase of theta phase consistency among later remembered trials, suggesting that the probability of successful encoding is higher for specific phases of the theta cycle.

**Figure 7.**
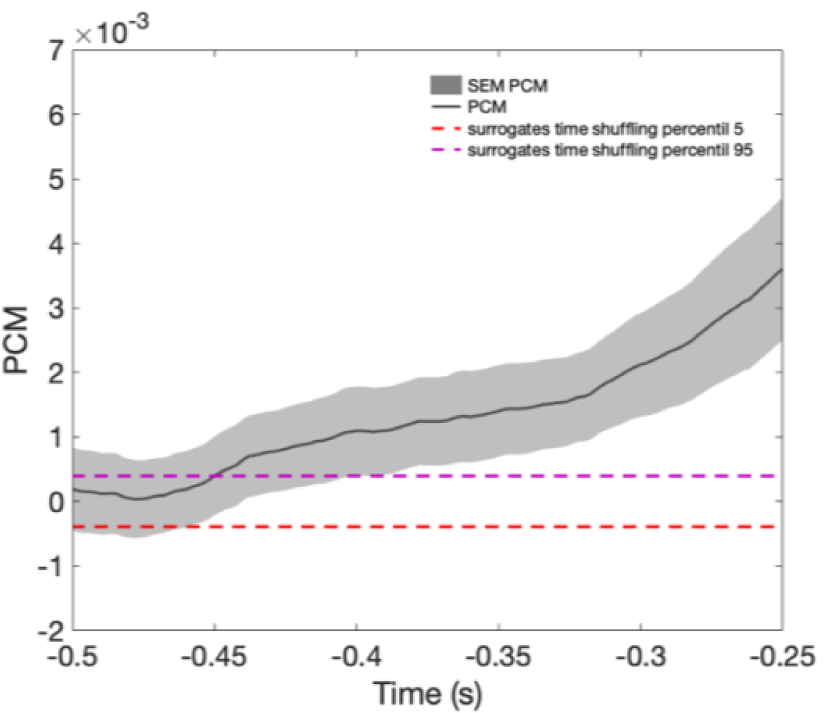
Phase concentration measure and surrogate distribution. To assess whether the surrogate distribution used was affected by a positive bias due to the concentration of hits trials, we used random distribution based on a different time window to calculate PCM surrogates, i.e., -0.8 to -0.5 s (because we know that no stimulus was presented there). Then, for each subject and electrode, labels for hit and miss trials were shuffled. For each trial, a random time point in the interval -0.8 to -0.5 was selected. Surrogate PCM was calculated based on these randomly sampled phases. The surrogate calculation was repeated 500 times for each subject and electrode and averaged across electrodes. 10.000 group surrogates were built (by sampling from individual surrogate distributions). We observed that no significant phase opposition was found (all PCM above 5% percentile surrogate distribution).

**Figure 8.**
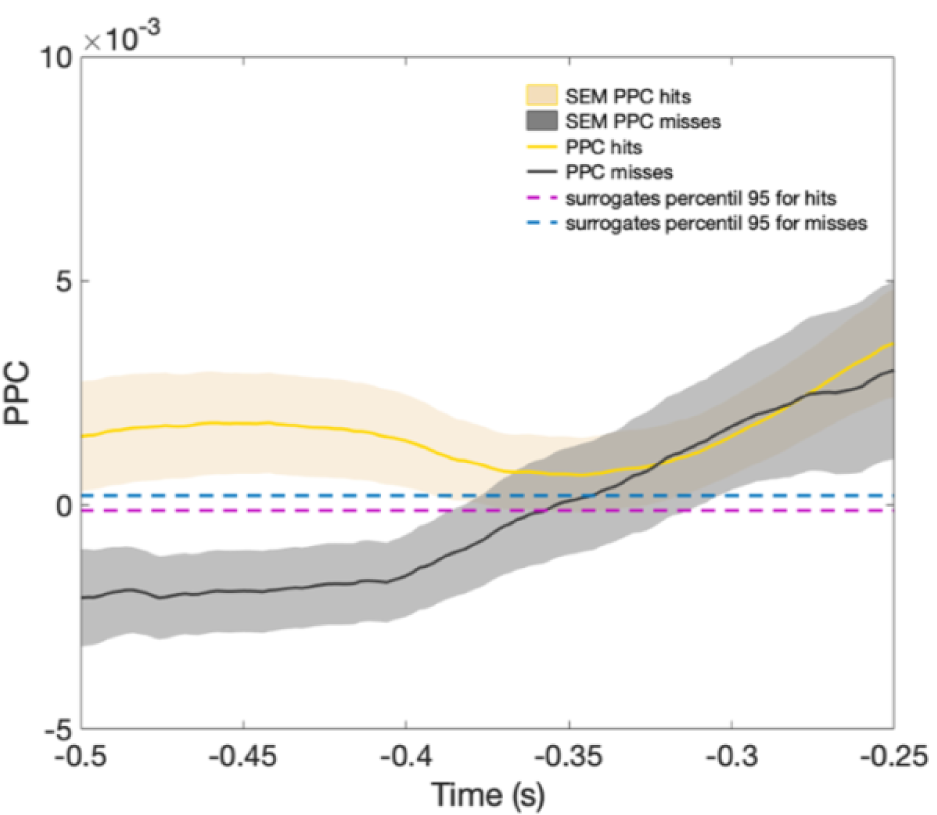
Phase concentration of hits and miss trials separately. The PPC for hit and miss trials was calculated separately and compared with the percentile 95 of a surrogate distribution. According to surrogate testing, hit phases are always concentrated (all p-values <0.0014, i.e., 251 consecutive timepoints), whereas, miss phases are concentrated in the time window from -0.356 to 0 s relative to stimulus onset.

## 6. Discussion

Although with mixed evidence, several studies have demonstrated a link between the phase of ongoing neural oscillations and behavioral outcomes, especially in perception and attention tasks (Busch et al. 2009; Mathewson et al. 2009; VanRullen et al. 2011). Conversely, a handful of studies have reported null results regarding this link (Ruzzoli et al. 2019; Bompas et al. 2015; Busch and VanRullen 2010; Benwell et al. 2017). The interpretation of these phase-behavior correlation results capitalizes on the idea that brain oscillations reflect fluctuations in neural excitability and the coordinated action of neural populations across different brain regions. Here, we applied the same logic to investigate whether the phase of ongoing activity at stimulus onset, has an impact on later memory performance. The question is whether fluctuations in prestimulus brain states, as reflected by oscillatory dynamics, will have an impact on memory formation and consequently, on subsequent recognition of the stimulus. We have obtained some positive evidence, after post-hoc analyses.

In the first experiment, we collected behavioral data from 30 healthy participants performing a visual paired-associates episodic memory task. Before each to-be-remembered image pair, an audio-visual reset cue was presented to induce modulation in cortical excitability by phase resetting the ongoing oscillatory activity (Lakatos et al. 2009; Fiebelkorn et al. 2011; Daitch et al. 2013). We measured later memory recognition performance as a function of the time-lag between the audio-visual reset cue and the presentation of the image pair to be encoded (randomly varied within an interval between 0 and 1000 ms, in steps of 16 ms). The data showed that subsequent memory performance for the associated pairs fluctuated periodically at ∼4 Hz as a function of the time lag from the audio-visual reset cue to encoding. Although the effect did not survive multiple comparisons correction for the number of frequencies included in the analysis, it suggests that theta-band oscillatory activity prestimulus may modulate the encoding. Consistent with previous findings, this periodic fluctuation could be attributed to a phase resetting of the ongoing oscillatory signal in the theta band that would be functionally relevant for encoding (Rizzuto et al. 2003; Fiebelkorn et al. 2011; Daitch et al. 2013; Fiebelkorn et al. 2018). However, the behavioral data in this experiment does not provide a direct measure of the phase of ongoing brain activity; therefore, we could only assume the phase resetting as the most likely explanation of the possible modulation. Because of this, and the lack of significance after correcting for multiple comparisons, we consider that any interpretation derived from this result must be cautious.

In a second experiment, we used EEG to investigate more directly the role of ongoing low-frequency brain oscillations prior to the to-be-encoded stimulus. Specifically, we focused on the phase and amplitude of frontotemporal theta and occipitoparietal alpha oscillations as a function of trial to trial successful or unsuccessful recognition. The results indicated that theta-phase differences between hits and misses in the prestimulus time-window predicted subsequent memory performance. Note that we pre-registered an analysis pipeline focusing on the phase of frontotemporal theta and occipitoparietal alpha that returned null results. After ascertaining that the ROI and the Phase Opposition Sum method (POS) (subsampling of trials with POS) were probably sub-optimal, we ran a post-hoc phase analysis based on the Phase Consistency Metric (PCM). This phase analysis was guided by a more precise estimation of the ROI and the individual frequency parameters based on post-stimulus theta power effects (i.e., increment after successfully remembered items). Following this data-driven approach, we observed significant prestimulus phase subsequent memory effects in a cluster of left frontal electrodes, suggesting a relationship between the phase of ongoing theta oscillations before stimulus onset, and later memory performance. Importantly, in the time window were the effect was observed, we only found phase concentration for hits, suggesting that there is a particular phase which favours encoding, and that the modulation observed in the hit rate relates exclusively to the hits concentration. This was not observed for miss trials.

The significant theta power effect, which was the base for the data-driven phase analysis, was a replication of a well-known SME consisting of a theta increase in the peri-stimulus period with similar topographic distribution as in previous studies (Osipova et al. 2006; White et al. 2013; Long et al. 2014; Khader et al. 2010; Guderian et al. 2009; Klimesch and Doppelmayr 1996). In particular—for each participant—the theta power effect appeared frontally after stimulus presentation and spread toward centro-posterior areas at increasing latencies. We may argue that the effects of phase and power are analytically independent since they were found at different times with respect to the stimulus onset. Increases in theta power appeared around (e.g., immediately before and during) stimulus presentation, while phase effects in the theta band predictive of later recognition were observed long before the stimulus, in the –500 to –442 ms, (as well as –134 to 82 ms) time window with respect to stimulus onset.

Most of the studies looking at prestimulus oscillatory effects in subsequent memory performance have used a prestimulus central orientation/fixation cue signaling the impending appearance of the to-be-encoded stimulus (Haque et al. 2015; Guderian et al. 2009; Otten et al. 2010). Thus, the increases in memory performance are mainly attributed to active anticipatory states. A substantial difference, at least with our second experiment, is that we focused on prestimulus ongoing oscillations; therefore, we did not use a prestimulus informative cue. In the first experiment, one could argue that the cue was not particularly time-informative (especially within the time window of relevance, 0-500 ms after the cue), although this could be more controversial. According to prior literature, in the absence of anticipatory states, i.e., under ongoing theta fluctuations, evidence from animal studies has shown an enhancement in the learning rate when the stimulus is presented during a specific hippocampal theta phase (Seager et al. 2002). The relevance of theta phase to the encoding of new information has been well-established through studies *in* vitro and in rodents and further implemented in leading theoretical models of memory (Huerta and Lisman 1995; Hyman et al. 2003; Hasselmo et al. 2002; Hasselmo 2005). Hippocampal theta is thought to be induced in cortical areas via hippocampal-cortical feedback connections, gating synaptic plasticity. In turn, the induction of long-term potentiation (LTP) is dependent on the phase of theta rhythm; whereas LTP preferentially occurs on the positive phase of the theta cycle, long-term depression occurs at opposing phases (Pavlides et al. 1988; Fell and Axmacher 2011).

Another possible interpretation for the role of the prestimulus phase in the modulation of later recognition success is that it may reflect attentional mechanisms. Indeed, attention orienting is known to impact memory encoding (Chun and Turk-Browne 2007). The idea is that the recruitment of frontal regions promotes encoding processes by top-down modulation of posterior occipitoparietal regions. In particular, frontal regions may contribute by selecting goal-relevant information and binding pieces of information (Gazzaley and Nobre 2012; Blumenfeld and Ranganath 2007). Thus, the fluctuations that have been observed may reflect cyclic changes in preparation for optimal stimulus processing (Sekuler and Kahana 2007; Gazzaley and Nobre 2012). Along these lines, Busch and colleagues (Busch and VanRullen 2010) showed that detection performance was improved by attention and fluctuated over time, along with the phase of spontaneous theta oscillations, before stimulus onset. However, the evidence against attentional mechanisms arises from studies suggesting that prestimulus attentional effects are correlated with decreases in occipitoparietal alpha (Thut et al. 2006; O’Connell et al. 2009; Mazaheri et al. 2009). In our study, we did not find significant prestimulus differences in alpha power, and our experimental design does not allow us to disentangle the effects of the induced anticipatory states from the impact of attentional processes.

Recognition and free recall are two memory processes whose performance could differ on storage, recovery operations, or some combination of both (Atkinson and Shiffrin 1968; Kahana 2012). In fact, using intracranial EEG, Merkow et al. (2014) showed that selective hippocampal prestimulus theta activity was associated with better subsequent recognition, but not with subsequent recall. Their results suggest that hippocampal prestimulus theta power increases preferentially promote the encoding of item information rather than the associative information of the item with the self-generated cues necessary for retrieval. Results from our analysis support the notion that prestimulus theta oscillations underlie mnemonic processes that favor later performance in recognition paradigms, but may not generalize to other memory paradigms, such as the free or cued recall.

One limitation of the present study is that the initial selection of the regions of interest was possibly not optimal. We initially divided the whole electrode set into two large clusters (anterior and posterior) that were too broad. We are aware that besides characterizing the functional significance of the observed effects in phase and amplitude, it is relevant to identify the brain areas that play a role in the observed effects. Although this is challenging to do with EEG, through additional analyses, we redefined the ROI as the set of electrodes displaying post-stimulus increases in theta power (putatively) after stimulus onset in our data, which replicate a relatively well known pattern (Osipova et al. 2006; Sederberg et al. 2003; White et al. 2013; Clouter et al. 2017; Khader et al. 2010; Guderian et al. 2009).

Another possible limitation of the present study was the choice of a suboptimal method for phase opposition analysis. The disproportion between correct trials and misses (as performance was well above chance level) resulted in a decrease of the sensitivity of the POS index (VanRullen, 2016). We found that our initial strategy to circumvent this problem, by equalizing the number of trials among conditions using subsampling, reduced the sensitivity of phase opposition measures more than anticipated (VanRullen 2016; Zoefel et al. 2019). Based on post-hoc simulations, we found that the decrease in the number of trials had a stronger impact than the imbalance itself. However, it remains an open question to fully understand how the combination of these two parameters (number of trials and balance in number of trials) affects the sensitivity of the phase opposition measures, something that is beyond the scope of this paper. In our follow-up exploratory analyses, we decided to use PCM, a measure that is less affected by trial imbalance, in order to use all available trials.

In summary, despite further confirmation will undoubtedly be needed, the principle finding to emerge from this study is that the spatiotemporal pattern of brain activity preceding the stimulus onset, measured with EEG, can predict behavioral performance in a memory recognition task. This provides further evidence that the state of neural activity preceding stimulus presentation has an impact on the subsequent processing, extending prior results in perceptual and attentional tasks. In the particular case of the memory task used here, the relevant spatiotemporal pattern is characterized by theta-band fluctuations and increases in phase consistency among hits reflected in the left frontal scalp. These novel insights highlight the role of theta phase in cortical oscillations at encoding and support episodic memory models linking behavioral data to phasic properties of theta rhythm (Hasselmo et al. 2002; Hasselmo 2005).

## 8. Acknowledgements

We thank Dr. Luis Fuentemilla and Dr. Pau Alexander Packard for their comments to an early version of the manuscript. We also thank the subjects for their participation in our study.

## 10. CRediT authorship contribution statement

**Josephine Cruzat:** Conceptualization; Data curation; Formal analysis; Investigation; Methodology; Visualization; Writing - original draft. **Mireia Torralba:** Conceptualization; Formal analysis; Investigation; Methodology; Software; Supervision; Validation; Visualization; Writing - review & editing. **Manuela Ruzzoli:** Conceptualization; Investigation; Funding acquisition; Methodology; Supervision; Writing - review & editing. **Alba Fernández:** Data curation. **Gustavo Deco:** Conceptualization; Methodology; Supervision; **Salvador Soto-Faraco:** Conceptualization; Funding acquisition; Investigation; Methodology; Supervision; Writing - review & editing.

## Supplementary Material

**Supplemental Table S1:** Demographic characteristics and performance for each participant in Experiment 1. Excluded participants are marked in red.

| Participant | Gender | Age | Handedness | Hit Rate (%) | False Alarm Rate (%) |
| --- | --- | --- | --- | --- | --- |
| 1 | F | 22 | R | 43,75 | 11,11 |
| 2 | F | 21 | R | 89,11 | 6,47 |
| 3 | M | 21 | L | 74,22 | 4,91 |
| 4 | F | 22 | R | 60,27 | 1,46 |
| 5 | F | 20 | L | 84,13 | 6,82 |
| 6 | F | 21 | R | 49,94 | 16,99 |
| 7 | F | 24 | R | 63,05 | 1,23 |
| 8 | F | 20 | R | 38,49 | 11,07 |
| 9 | F | 25 | R | 95,36 | 5,65 |
| 10 | M | 21 | R | 64,29 | 2,90 |
| 11 | F | 42 | R | 71,85 | 3,35 |
| 12 | F | 21 | R | 73,64 | 1,55 |
| 13 | F | 23 | R | 66,78 | 4,59 |
| 14 | M | 22 | R | 58,49 | 6,62 |
| 15 | F | 20 | R | 81,94 | 4,41 |
| 16 | F | 19 | R | 68,48 | 11,34 |
| 17 | F | 20 | R | 49,76 | 8,39 |
| 18 | F | 20 | R | 68,67 | 1,06 |
| 19 | F | 21 | R | 76,67 | 8,91 |
| 20 | M | 21 | R | 85,23 | 9,15 |
| 21 | M | 24 | R | 55,05 | 7,34 |
| 22 | F | 20 | R | 67,14 | 3,32 |
| 23 | M | 27 | R | 45,00 | 12,74 |
| 24 | M | 23 | R | 66,18 | 2,27 |
| 25 | F | 21 | R | 76,62 | 3,03 |
| 26 | M | 26 | L | 47,23 | 9,30 |
| 27 | M | 20 | L | 79,90 | 3,52 |
| 28 | F | 21 | R | 52,58 | 7,91 |
| 29 | F | 32 | R | 67,21 | 4,15 |
| 30 | F | 31 | R | 58,60 | 7,10 |
| 31 | F | 22 | R | 66,54 | 4,47 |
| 32 | F | 24 | R | 87,79 | 8,33 |
| 33 | M | 23 | R | 65,57 | 12,15 |
| 34 | F | 22 | R | 91,07 | 3,23 |
| 35 | F | 21 | R | 68,76 | 11,30 |
| 36 | F | 24 | R | 76,33 | 1,83 |
| 37 | F | 32 | R | 54,18 | 10,94 |
| 38 | M | 26 | L | 64,07 | 3,04 |

**Supplemental Table S2:** Demographic characteristics and performance for each participant in Experiment 2. Excluded participants are marked in red.

| Participant | Gender | Age | Handedness | Hit Rate (%) | False Alarm Rate (%) |
| --- | --- | --- | --- | --- | --- |
| 1 | F | 22 | R | 58,31 | 9,03 |
| 2 | M | 26 | R | 58,91 | 10,25 |
| 3 | M | 21 | R | 75,53 | 10,39 |
| 4 | F | 21 | R | 56,80 | 13,33 |
| 5 | **M** | **29** | **R** | **45,63** | **27,76** |
| 6 | **F** | **23** | **R** | **87,67** | **4,38** |
| 7 | F | 22 | R | 46,12 | 6,51 |
| 8 | **F** | **23** | **R** | **78,25** | **2,49** |
| 9 | M | 23 | R | 70,52 | 5,00 |
| 10 | F | 23 | L | 61,37 | 8,51 |
| 11 | M | 24 | R | 52,73 | 11,66 |
| 12 | M | 24 | R | 63,72 | 0,00 |
| 13 | M | 26 | R | 66,98 | 6,57 |
| 14 | F | 29 | R | 56,40 | 7,80 |
| 15 | F | 26 | R | 71,46 | 10,80 |
| 16 | F | 20 | R | 68,27 | 3,82 |
| 17 | M | 22 | R | 64,59 | 9,09 |
| 18 | **M** | **19** | **R** | **77,62** | **6,34** |
| 19 | **M** | **27** | **R** | **76,33** | **6,23** |
| 20 | **F** | **19** | **R** | **77,14** | **4,93** |
| 21 | F | 23 | R | 69,17 | 4,45 |
| 22 | M | 20 | R | 60,29 | 4,90 |
| 23 | F | 23 | R | 76,56 | 1,40 |
| 24 | M | 18 | R | 70,26 | 10,10 |
| 25 | M | 19 | L | 70,12 | 6,81 |
| 26 | M | 25 | R | 66,51 | 3,11 |
| 27 | M | 20 | R | 60,14 | 11,23 |
| 28 | F | 22 | L | 63,36 | 7,12 |
| 29 | F | 21 | R | 69,09 | 1,08 |
| 30 | F | 23 | L | 60,43 | 10,10 |
| 31 | F | 20 | R | 72,60 | 7,22 |
| 32 | F | 23 | R | 67,29 | 5,38 |
| 33 | **M** | **22** | **R** | **74,54** | **4,04** |
| 34 | M | 20 | L | 60,19 | 4,73 |
| 35 | M | 25 | R | 57,41 | 10,39 |
| 36 | M | 26 | R | 67,76 | 12,68 |
| 37 | F | 31 | R | 61,56 | 10,36 |

**Supplemental Figure S1:**
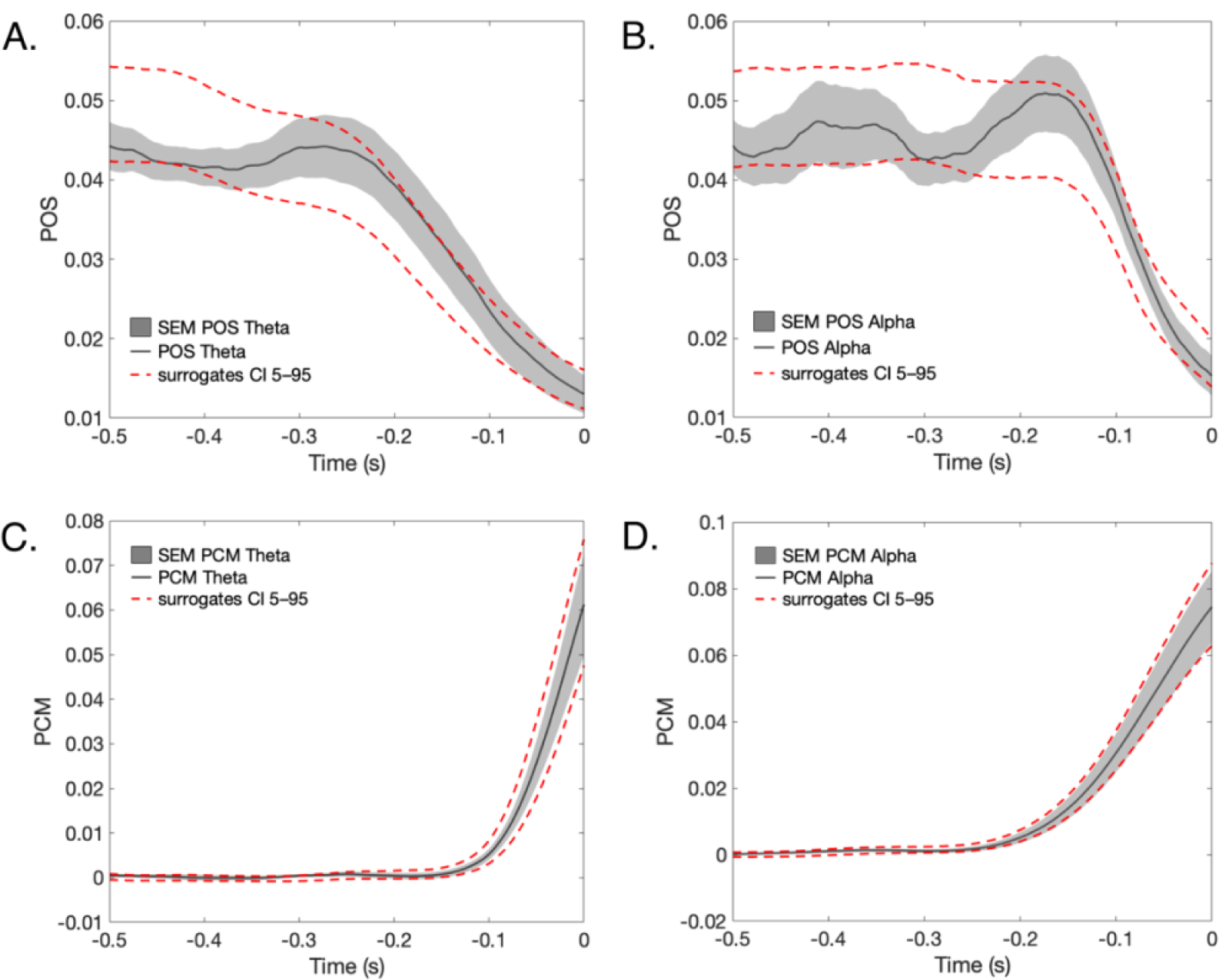
Phase consistency metrics. POS analysis showed no significant phase opposition in either (A) Theta in frontotemporal electrodes (max 0.044 ± 0.017, t=-500 ms), or (B) Alpha in occipitoparietal electrodes (max 0.0509 ± 0.026, t=-174 ms). Similar to the POS, the PCM showed null results for (C) Theta in frontotemporal electrodes (min 0.00013 ± 0.0022, t=-500 ms), or Alpha in occipitoparietal electrodes (min: 0.000087 ± 0.0027, t=-350 ms).

**Supplemental Figure S2:**
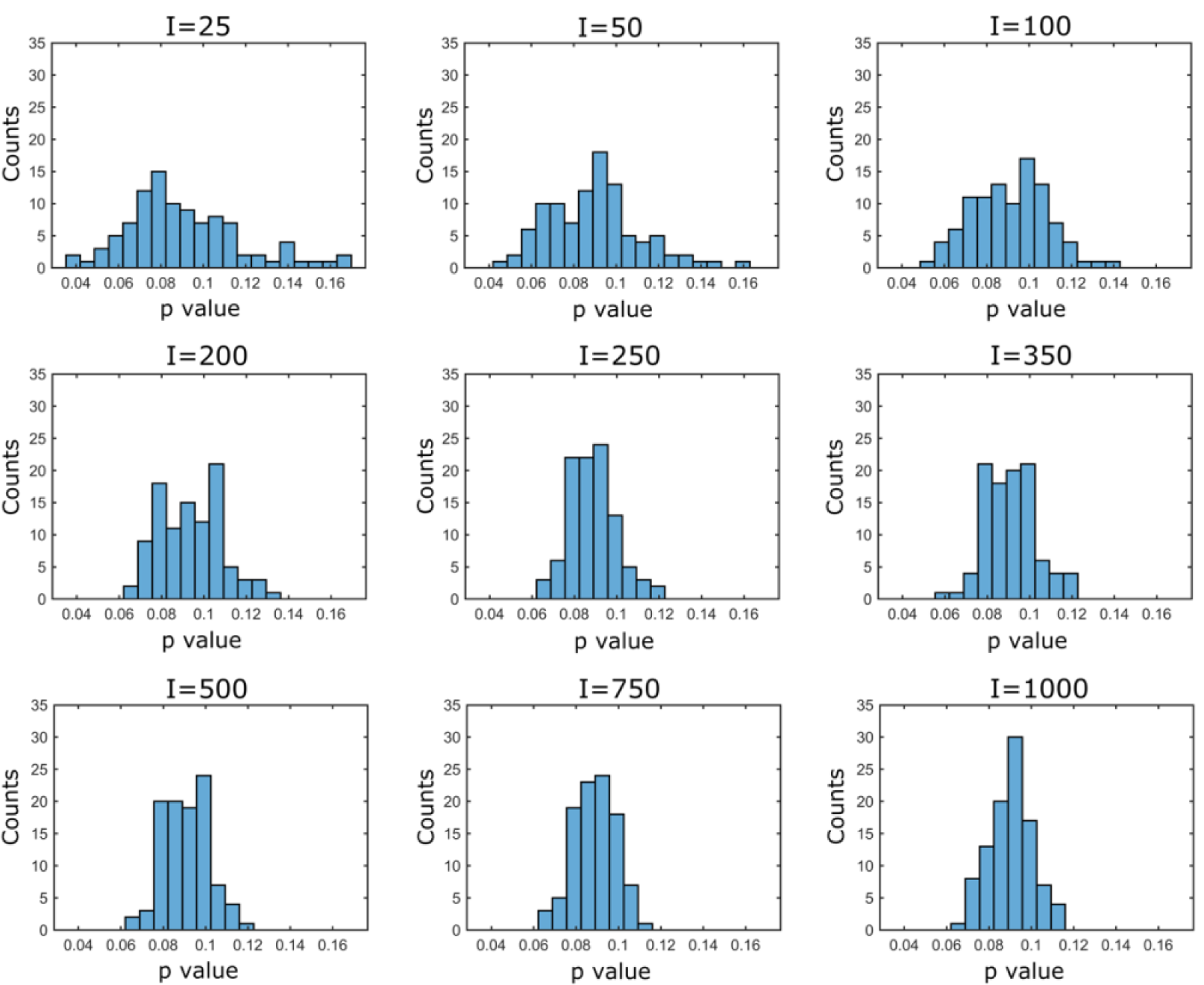
Assessing the stability of the POS measure. Histograms of p-values obtained for POS analysis for one electrode and time point in the theta frequency band, for different number of random (I) pickings per participant. Each histogram contains 100 values. As can be seen, the p-values obtained were not stable: at the maximum number of iterations, the p-values varied in the interval 0.06 to 0.12

**Supplemental Figure S3:**
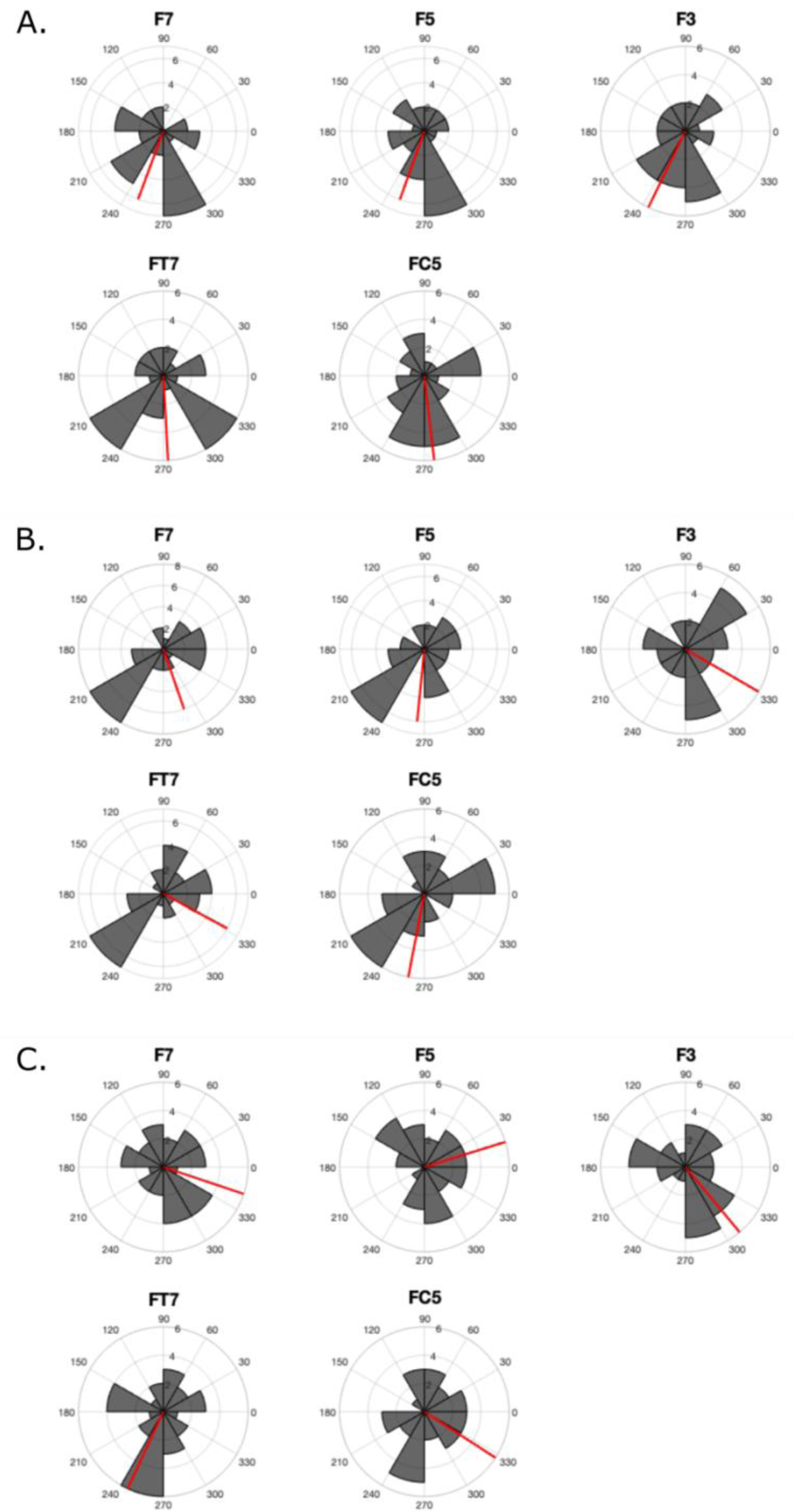
Differences in (A) mean hit phases (B) mean miss phases and (c) difference between hit and miss phases, for all subjects, in the most significant and positive timepoint of PPC_Hits_ in the time window where PPC_Miss_ is not significant (t=-442 ms), for electrodes in the frontal areas. As can be seen, phases for hits trials were close to opposition for some of the electrodes, as compared to phases of miss trials suggesting an optimal phase for memory encoding.

**Supplemental Figure S4:**
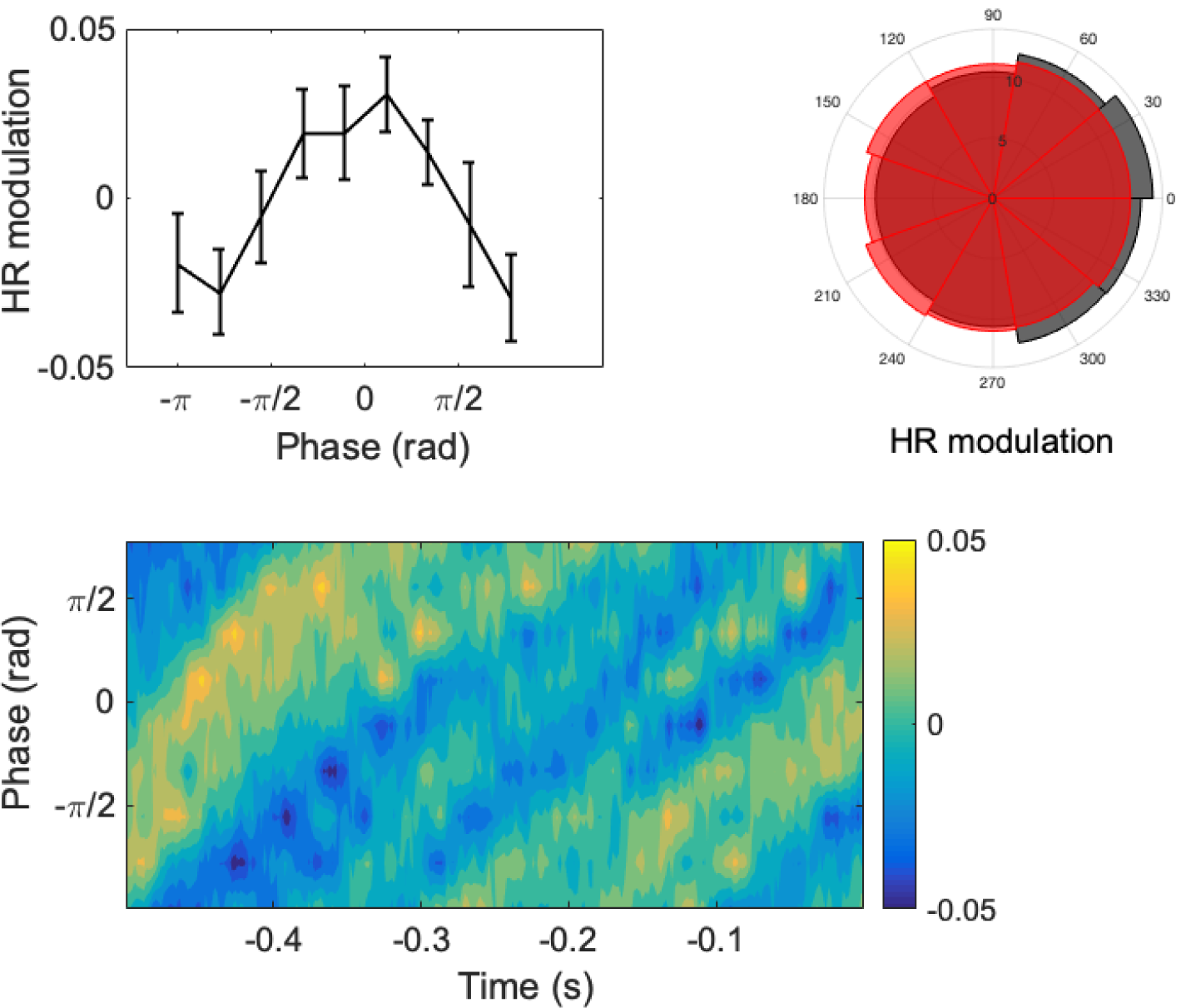
Phase distributions for all trials at the electrode with the lower p-value and most positive PPC for hits (electrode F7)(∼442 ms). Panel A shows variations in hit rate (HR) with respect to mean HR at -442 ms (mean HR=0.647, thus, 0.05 corresponds to a 7.7% change in HR). Panel B shows percentage of total hits (grey) and misses (red) at each phase bin at -442 ms. Panel C shows HR modulation as a function of time. In order to reduce interparticipant variability, for each subject, phases were aligned to the mean hit phase at -442 ms.

